# Heterogeneous ‘cell types’ can improve performance of deep neural networks

**DOI:** 10.1101/2021.06.21.449346

**Authors:** Briar Doty, Stefan Mihalas, Anton Arkhipov, Alex Piet

## Abstract

Deep convolutional neural networks (CNNs) are powerful computational tools for a large variety of tasks (Goodfellow, 2016). Their architecture, composed of layers of repeated identical neural units, draws inspiration from visual neuroscience. However, biological circuits contain a myriad of additional details and complexity not translated to CNNs, including diverse neural cell types (Tasic, 2018). Many possible roles for neural cell types have been proposed, including: learning, stabilizing excitation and inhibition, and diverse normalization (Marblestone, 2016; Gouwens, 2019). Here we investigate whether neural cell types, instantiated as diverse activation functions in CNNs, can assist in the feed-forward computational abilities of neural circuits. Our heterogeneous cell type networks mix multiple activation functions within each activation layer. We assess the value of mixed activation functions by comparing image classification performance to that of homogeneous control networks with only one activation function per network. We observe that mixing activation functions can improve the image classification abilities of CNNs. Importantly, we find larger improvements when the activation functions are more diverse, and in more constrained networks. Our results suggest a feed-forward computational role for diverse cell types in biological circuits. Additionally, our results open new avenues for the development of more powerful CNNs.

## Introduction

Deep convolutional neural networks (CNNs) draw architectural inspiration from visual neuroscience. CNNs contain many processing units that aim to emulate the role of neurons in the mammalian visual system, a series of iterative processing steps (Goodfellow, 2016, Hubel & Wiesel, 1959; Hubel & Wiesel, 1962). Recently performance of CNNs has rapidly improved across a wide range of computational tasks, particularly visual object recognition and classification (LeCun et. al., 2015). CNNs work by developing highly nonlinear representations of inputs across repeated layers of processing units. These representations are learned through training by updating the connections, or weights, between units in successive layers. Types of processing layers include convolutional layers, which convolve input with learned spatial filters, activation layers which include a nonlinear activation function, and max-pooling layers. Typically, CNNs units are uniform across each layer within a network. Network architectures—meaning the number, size, type, and arrangement of processing unit layers—are often manually designed through iterative experimentation, although automated search techniques are being actively developed (Zoph & Le, 2016). Individual convolutional features and weights are determined through iterative training, not architectural design.

Biological circuits, however, contain diverse cell types at each location in the visual hierarchy. Classification of neural cell types dates back to the pioneering work of Ramon y Cajal, who described in fine detail elaborate neural morphology (Llinas, 2003). Recent studies have classified neurons by their location in the brain, morphology, gene transcription, and electrical properties (Tasic, 2018; Teeter, 2018; Gouwens, 2019; Gouwens, 2020). The biological function of these diverse neural cell types is not yet fully understood, but is an active area of research (Burnham, 2021; Zeldenrust, 2021; Perez-Nieves, 2021). Many possible roles for neural cell types have been proposed, including: learning, stabilizing excitation and inhibition, and diverse normalization (Marblestone, 2016; Gouwens, 2019). Recently several studies have begun to investigate computational properties of networks with heterogeneous cell types, instantiated in different ways such as excitatory vs inhibitory (Cornford, 2020), synapses (Burnham, 2021), connectivity (Stöckl, 2021), and intrinsic dynamics (Padmanabhan, 2010; Gjorgjieva, 2016; Duarte, 2019; Perez-Nieves, 2021; Zeldenrust, 2021). Broadly, these studies find computational benefits from adding heterogeneity to neurons. Burnham et al, 2021 and Perez-Nieves et al, 2021 investigated adding heterogeneous synaptic and membrane timescales, finding improved performance on standardized sequential datasets. Stöckl et al, 2021 constructed networks with cell type specific connectivity rules, and found improved performance with reduced number of neurons. Zeldenrust et al, 2021 analytically derived a class of spiking network models that optimally track time-varying inputs, resulting in networks with diverse internal dynamics. We build on these results by instantiating cell type diversity as mixed activation functions, and explore the possible role for neural heterogeneity in feed-forward computation in CNNs performing image classification.

One basic classification of neurons uses their electrical input-output relationships characterizing them electrophysiologically based on features of their *f-I* curves—nonlinear functions that map input current to neuronal firing rate (Izhikevich, 2007). In the deep learning context, neurons are reduced to units represented by a single, 1-dimensional quantity: their activation function. Activation functions are typically nonlinear functions that transform the weighted sum of a unit’s input to an output value. Taking advantage of this similarity between activation functions and *f-I* curves, we can designate different CNN unit cell types by applying mixed activation functions as an analog for electrophysiological classification. Applying these constraints in the deep learning context presents an opportunity to use CNNs as a model for studying the role that heterogeneous cell types might play in feed-forward sensory coding. Here we investigate the effects of adding mixed activation functions as an analog for cell types to an existing CNN architecture. To constrain our investigation we did not explore heterogeneous connectivity (Stöckl, 2021), synapses (Burnham, 2021), timescales (Burnham, 2021 and Perez-Nieves, 2021), or spike-timing (Zeldenrust, 2021). We seek to characterize the network response to the addition of cell types in terms of classification accuracy, learning, and the network’s internal representation of the input space. We find that mixed activation functions can improve image classification compared to control homogeneous networks, and that the benefit of mixed activation functions is larger in more constrained networks. Finally, we find that internal representations in mixed networks differ from the control networks.

## Results

### Mixed activation functions on an image classification task

To investigate the role of heterogeneous cell types in feed-forward processing, we implemented a programmatic framework that generates CNN instances containing heterogeneous configurations of activation functions, then trains them on an image classification task. We adopted the CIFAR-10 image set as our primary classification benchmark. This imageset contains 10 image classes with 6000 exemplars in each class (Krizhevsky, 2009). As a standard network testbed we used the popular, relatively simple, VGG11 architecture (Simonyan & Zisserman, 2014). VGG11 contains 11 composite layers including 8 convolutional layers and 3 fully connected layers. We started experimentation with VGG11 because its uniform architecture across layers allows for straightforward analysis and modification, its popularity means its performance is well-studied on popular datasets, and its depth offers the ability to attain state of the art classification accuracy.

We replaced the homogeneous activation functions (Figure 1a) in VGG11 to introduce heterogeneity (Figure 1b). In particular, each mixed network contains two activation functions within every activation layer and fully connected layer. We refer to a network configuration that mixes activation functions A and B with a name of the form “mixed A-B” (Figure 1b), and a homogeneous control network that uses only A or B as “control A” or “control B” (Figure 1a). Note that our mixed networks contain the same number of units, connections, and learnable parameters as the control networks. For each mixed or control configuration, we trained 10 initializations of the same network architecture with random starting weights. This approach allows us to statistically assess whether mixed activation functions improve image classification ability. For example, comparing the control networks Swish(7.5), and PTanh(1) (Figure 1c), with the corresponding mixed network Swish(7.5)-PTanh(1), we find the mixed network outperforms both the linear average of the two control networks, and each control network individually (Figure 1d).

**Figure 1.**
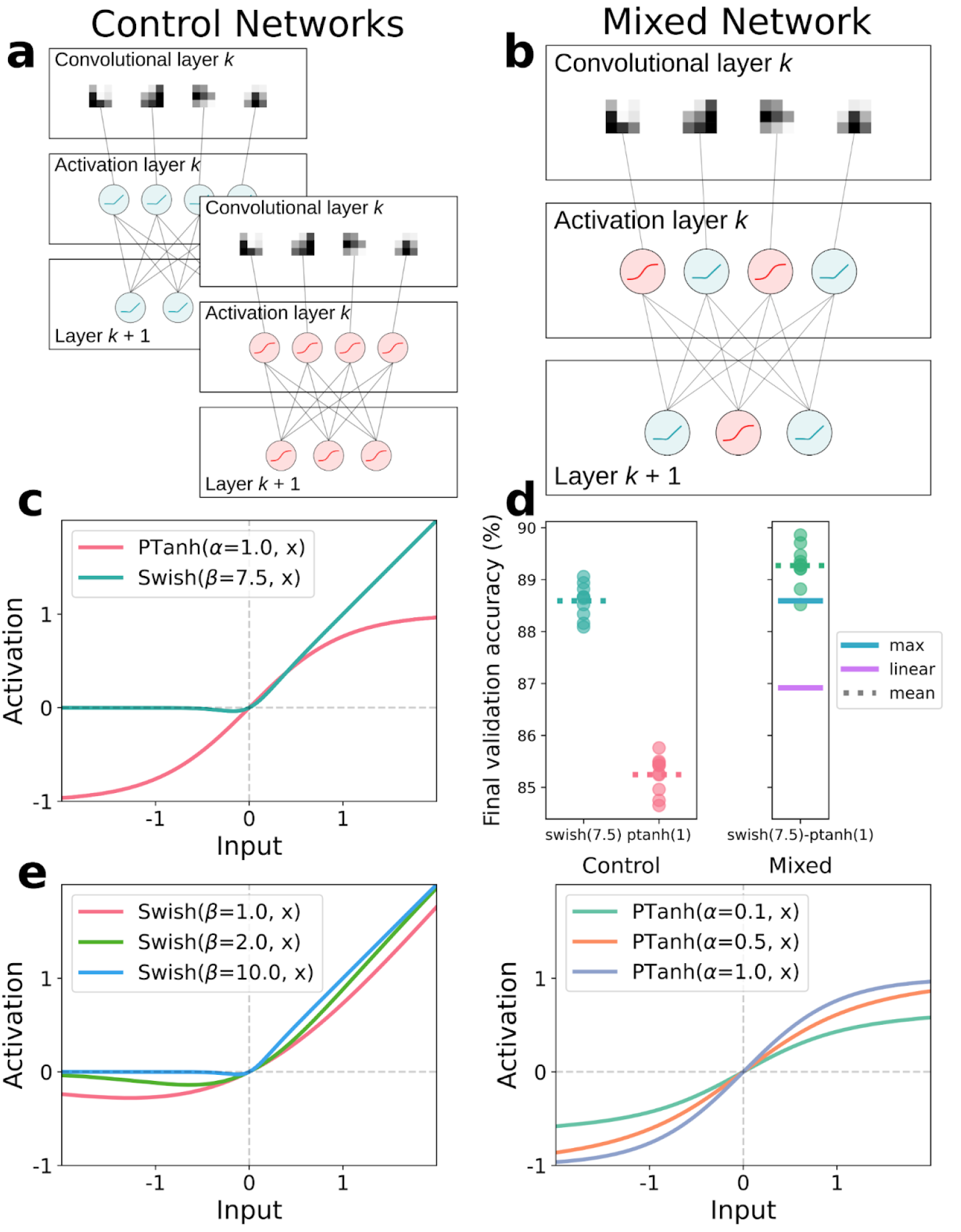
**(a)** Homogeneous control networks corresponding to the mixed network in (b). Each control network contains only one activation function across all units. **(b)** Illustration of mixed activation functions within a CNN. PTanh and Swish units are intermixed within each activation and fully connected layer in the network. **(c)** PTanh and Swish nonlinear activation functions. PTanh is saturating, while Swish has rectification properties. **(d)** Comparison of final classification accuracies for a heterogeneous configuration of VGG11 and its corresponding control networks. For each network type, each dot represents the final accuracy of a single random initialization, and the dashed line is the mean accuracy across initializations. The max prediction is the greater of the two control cases, while the linear prediction is the average of the two control cases. Here the mixed network outperforms both predictions. **(e)** Examples of Swish and PTanh nonlinearities over a range of parameter values.

### Search space

To keep the scope of the investigation practical, we defined a search space that balances dynamic range of effect with the feasibility of exploration. The dimensions of this space include the set of candidate activation functions and their associated parameters, the ten targetable activation layers in VGG11, and the set of locations within the targeted layers at which activation functions can be applied. Since an exhaustive exploration of all possible permutations of even a small set of activation functions in a single layer is not feasible, we set out to target all activation layers with systematic cell type arrangements that offer a good sampling of the overall search space. Our results presented here used a 1:1 ratio of two activation functions within each layer of each network.

### Families of parametric activation functions

The choice of activation functions in artificial deep networks is an active area of research (Ramachandran et. al., 2017). Different activation functions have been historically used in deep learning with different motivations (Goodfellow et. al., 2016). We separated activation functions into different families that exhibit the qualitatively different behaviors of rectification and saturation (Figure 1c). Commonly used members of each family are thought to offer different advantages in deep neural networks. For example, rectifying functions like ReLU are often piecewise linear, and are thought to help the network develop a sparse internal representation of the input space. Saturating activation functions like Tanh operate with bounded outputs at input extremes. And a non-monotonic activation function like Swish offers behavior that is mostly rectifying, while maintaining continuous differentiability. We reasoned that mixed activation functions might offer computational benefits by combining the diverse advantages of different function families. We chose the parametric activation function Swish to represent the rectifying family (eq. 1), and the saturating activation function PTanh of our own design to represent the saturating family (eq. 2).

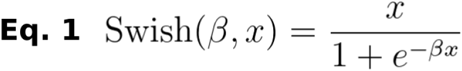

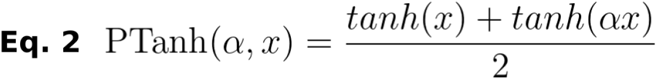

In addition to mixing across function families, we can evaluate the benefits of mixing within a function family by changing the function hyper-parameters ***β*** and α. Over a range of parameters, a single nonlinearity like Swish or PTanh can take a variety of forms (Figure 1e). PTanh has a parameter α that changes the curvature of the activation function such that the network cannot compensate by scaling the input weights. As the Swish parameter ***β*** increases, the function displays increasing rectification, becoming more similar to ReLU. Here we explore mixing cell types both “within” function families: the same function with a different parameter value, and “across” function families: different functions with potentially different parameter values.

### Accuracy relative to linear and max prediction

To evaluate the role of mixed activation functions, the performance of a given mixed network is compared to that of its corresponding homogeneous control networks using two metrics: the mixed network’s linear and max prediction (Figure 1d). The linear prediction expects the mixed network to perform as well as a linear combination of its control network accuracies. Our mixed networks use a 1:1 ratio of two activation functions, so the linear prediction is simply the average of the two control accuracies. For example, the control networks Swish(***β***=7.5) and PTanh(*α*=1) have mean final accuracies 88.59% and 85.24%, therefore the linear prediction for mixed network Swish(***β***=7.5)-PTanh(α=1) is 86.92%. Note here that the mean final accuracy for both the homogeneous control networks and the mixed networks is the average over 10 random initializations of each network type. The max prediction accounts for the possibility that a mixed network learns to compensate for the introduction of a population of units with a nonlinearity that performs poorly by adjusting its weights to “ignore” those units, thereby inaccurately inflating the mixed net’s performance with respect to its linear prediction. The max prediction is resistant to this effect. It expects the mixed network to perform as well as the best of its control network accuracies. For example, given the same control network accuracies presented above, the max prediction for mixed network Swish(7.5)-PTanh(1) is 88.59%. Figure 1d shows results for Swish(7.5)-PTanh(1), with mean final accuracy of 89.27%, outperforms both the maximum and linear prediction of its control networks. Our primary investigation aims to uncover under what conditions mixed activation function networks can outperform the max prediction of their control networks.

### Diverse heterogeneous networks outperform homogeneous control networks

Using the VGG11 architecture, we applied heterogeneous configurations within and across the parametric spaces of Swish and PTanh nonlinearities, then compared the final validation accuracies attained by each configuration on CIFAR-10 to the linear and max prediction based on each configuration’s corresponding homogeneous control networks. We observe that cross-family networks and within-family networks composed of diverse nonlinearities frequently beat their linear predictions (34 out of 35 cross-family and 21 out of 31 within-family using VGG11). However, when we hold the same network configurations to the higher standard of the max prediction (see example in Figure 1d), network performance relative to the control networks significantly depends on the choice of activation function parameters.

Using the max prediction benchmark, 6 out of 35 cross-family heterogeneous configurations impart an improvement in image classification accuracy to VGG11 that is statistically significant after adjusting for multiple comparisons (Figure 2a, note that this panel excludes networks containing nonlinearities Swish(***β***=0.1) and PTanh(α=0.01) for visual clarity). We refer to a network that beats its max prediction as exhibiting a positive response to mixed activation function. The largest positive response is that of Swish(7.5)-PTanh(1), which beats its max prediction by 0.7% on average. The magnitude of this effect corresponds to around 70 additional correct image classifications in CIFAR-10’s test set that neither control network configurations Swish(7.5) or PTanh(1) was able to correctly classify on average. The mean validation accuracy relative to max prediction for cross-family configurations of VGG11 is +0.04% (Figure 2b), and +0.79% relative to the linear prediction (Figure 2c). VGG11’s cross-family parameter landscape is relatively smooth, with a clear preference for higher Swish ***β*** values (Figure 2d). Specifically, we find configurations that mix PTanh(α) with Swish(***β***) functions using higher ***β*** values exhibit the largest increase in accuracy, and the highest final accuracy overall (Figure 2d). However, we see a less clear preference for PTanh’s α values. It is worth noting that as Swish’s ***β*** increases, the Swish function has stronger rectification and the activation profile more closely approximates that of ReLU (Figure 1e). These results demonstrate that heterogeneous activation functions can improve image classification.

Within-family heterogeneous configurations of VGG11 rarely outperform their max predictions (Figure 2b)—none of the 31 such configurations explored beat their max prediction with statistical significance. This demonstrates that the benefit of mixed activation functions may arise from utilizing the diverse characteristics of activation function families, such as saturation and rectification.

### Sticknet8—a reduced-parameter CNN

Between its filters, biases, and weights, VGG11 contains over 120 million trainable parameters. Additionally, since VGG11 has been demonstrated to achieve over 90% testing accuracy on CIFAR-10 (Fu 2019), there remains less than 10% left for potential improvement beyond that as a result of the addition of heterogeneous activation functions. Motivated by the factors above, we reasoned that more significant improvements may be readily observable in a network more constrained than VGG11. Therefore we set out to investigate the effects of adding mixed cell types to Sticknet8—a CNN which we describe here, based on the VGG11 architecture, but with approximately 1000 times fewer parameters. Sticknet8 joins the first 2 convolutional layers from VGG11 with VGG11’s final 3 fully-connected layers, while significantly reducing the number of parameters per layer. It preserves aspects of VGG11’s architecture including its 3×3 convolutional filters, its repeated groups of convolutional, activation, and pooling layers, and its relative layer-to-layer increases in parameters, but reduces the number of filters in the first convolutional layer from 64 to just 8. These changes result in a reduction from VGG11’s 128 million parameters down to Sticknet8’s 119,000. Using Sticknet8 as a base architecture, we repeated the experiments performed on VGG11 with the same set of heterogeneous network configurations.

**Figure 2.**
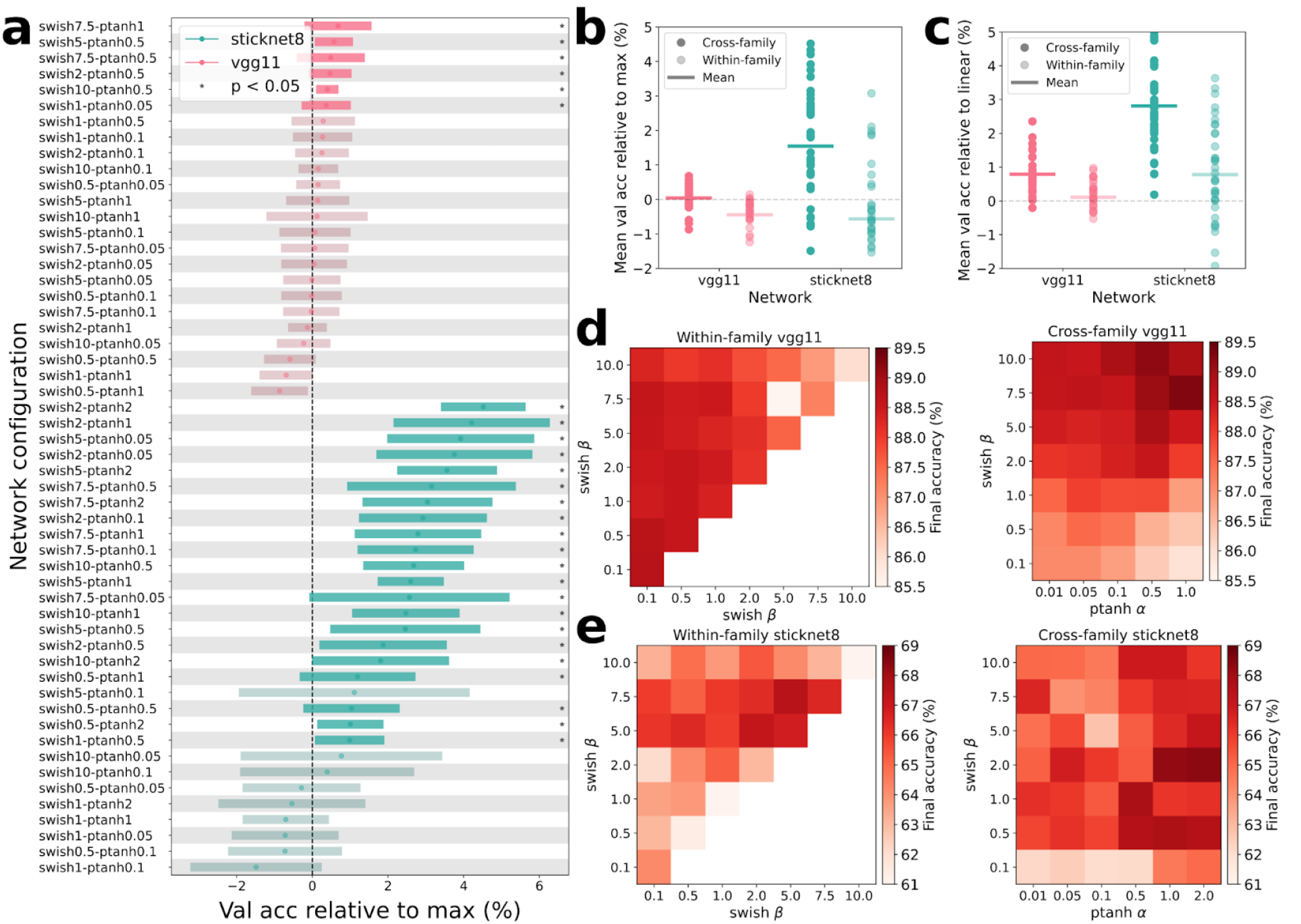
Mixed activation functions improve image classification performance. **(a)** Summary of mixed network accuracy relative to max prediction compared to control networks for VGG11 (red) and Sticknet8 (teal) architectures. Only cross-family networks are shown for clarity. Error bars represent a 95% confidence interval across 10 independently trained random initializations. Stars and darker shading indicate significant improvement of mixed network performance over control networks (one-sided t-test, p < 0.05 after Benjamini-Hochberg correction for multiple comparisons). Networks containing nonlinearities Swish(***β***=0.1) and PTanh(α=0.01) excluded for visual clarity (present in remaining panels). **(b)** Comparison of mean peak accuracy relative to max prediction for cross- and within-family mixed network configurations of VGG11 (only significant networks included). **(c)** Same as (b) using linear prediction. Notably, both cross- and within-family mixed networks beat their linear predictions on average for both network architectures. **(d)** Peak accuracy parameter landscapes for within-family Swish networks (left) and cross-family networks (right) for VGG11 architecture. **(e)** Same as (d) using Sticknet8 architecture.

### Sticknet8 offers more pronounced benefits of mixed activation functions

As expected, the CIFAR-10 final validation accuracies for Sticknet8 are generally lower than they are for the less constrained VGG11. The best configurations of Sticknet8 attain peak accuracies around 68% or lower (Figure 2e). However, the positive responses to the addition of cross-family mixed nonlinearities are both more frequent across explored network configurations (Figure 2a) and more pronounced within each configuration for Sticknet8 than for VGG11 (Figure 2b), supporting our expectation that improvements would be more readily observable in a parameter-constrained network that operates in a lower peak accuracy regime of CIFAR-10 than VGG11. Consistent with VGG11, the max prediction is a higher standard than the linear prediction, and cross-family networks outperform within-family networks. Compared to the linear prediction, 42 out of 42 cross-family, and 27 out of 36 within-family configurations outperform the linear prediction. Using the max prediction benchmark, 21 of the 42 cross-family configurations of Sticknet8 exhibit a positive response to the introduction of mixed nonlinearities (Figure 2a, note again some network configurations excluded from figure for visual clarity), and 3 out of 36 within-family configurations outperformed their max predictions.

While the effects in Sticknet8 hold over a larger landscape of parameter values, the performance is less sharply dependent on specific parameter values (Figure 2e). Interestingly, the best performing parameter combinations differ between Sticknet8 and VGG11, with Sticknet8 achieving the best performance with smaller values of Swish’s ***β*** compared to VGG11 (Figure 2d,e). Notably, several mixed configurations that negatively impacted the performance of VGG11 are among the top performing configurations of Sticknet8 (Figure 2a): Swish(2)-PTanh(1), Swish(5)-PTanh(0.05), and Swish(2)-PTanh(0.05). These three configurations beat their max predictions by almost 4%.

Our results with Sticknet8 demonstrate the computational advantages of mixed activation functions are more pronounced in a more constrained network, with mixed activation configurations beating their max predictions both more frequently and with larger magnitudes. As with VGG11, we observe the largest benefit from mixing across activation function families.

### Activation functions alter dimensionality of network responses

To gain insight into when and how mixed activation functions improve classification accuracy, we measured the dimensionality of network activations within each layer in response to a stratified subset of 500 CIFAR-10 images. We measured dimensionality using the participation ratio. The participation ratio measures the dimensionality as the square of the sum of the covariance matrix of unit activation eigenvalues over the sum of the squared eigenvalues (Recanatesi, 2019a; Recanatesi, 2019b; Gao, 2017; Rajan, 2010):

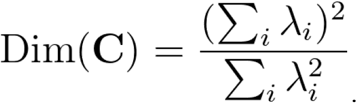

We adopted the participation ratio because it is simple to compute and interpret.

**Figure 3.**
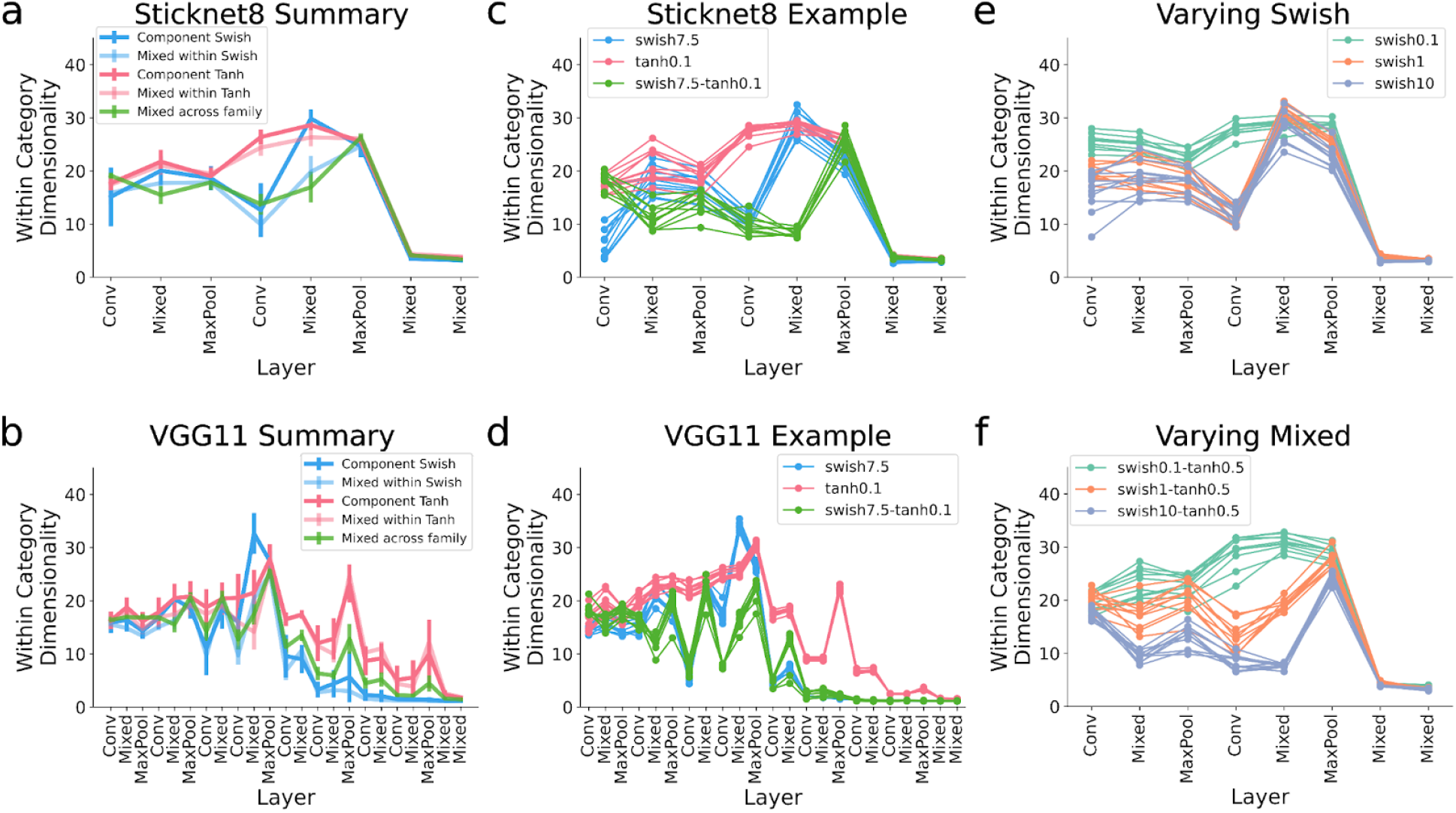
Different activation functions change the dimensionality of internal representations. **(a)** The average dimensionality of the response to each image category (within category dimensionality) for different classes of Sticknet8 networks (mean ± 95% CI). **(b)** Same as (a) for VGG11 networks. **(c)** Dimensionality across layers for an example Sticknet8 mixed network and its control networks (colors). Each network has 10 random initializations (lines). **(d)** Same as (c) for VGG11. **(e)** Dimensionality across three swish control networks with different parameter values. **(f)** Dimensionality across three cross-family mixed networks, varying the swish parameter, and fixing the tanh parameter.

We find in both Sticknet8 and VGG11 that changing the activation function dramatically alters the dimensionality of network activations within each layer (Figure 3a,b). Consistent with previous findings (Recanatesi, 2019a), in both Sticknet8 and VGG11 we find dimensionality generally increases moving deeper into the network before decreasing in the final layers (Figure 3a,b). Additionally, in both architectures, the dimensionality was largely consistent across the 10 initializations of each network configuration, with consistency increasing in deeper layers (Figure 3c,d). The relationship between activation functions and dimensionality within each layer appears complex. As a demonstration, Figure 3 shows the dimensionality of several example network configurations. Mixed networks generally displayed dimensionalities that differed from their control networks, and did not appear to follow a linear relationship (Figure 3c,d). The dimensionality responds nonlinearly to smooth variation of the activation function parameter (Figure 3e). In a mixed network configuration smoothly varying the parameter of one activation function in the presence of a constant second activation function resulted in large changes in dimensionality (Figure 3f). For example in Figure 3, comparing each Swish() control network in Figure 3e with the corresponding mixed network in Figure 3f, we find the layer with the largest dimensionality shifts to a deeper layer. In the case of Sticknet8, cross-family networks and within-family Swish networks—but not within-family PTanh networks—displayed smaller dimensionality throughout layers than control networks on average (Figure 3a). We did not observe this pattern in VGG11 (Figure 3b). This discrepancy may underlie the increased benefit of mixed activation functions in Sticknet8 compared to VGG11. In summary, we find that modifying activation functions can dramatically alter the internal representations of deep convolutional networks.

## Discussion

Our results show that the addition of heterogeneous activation functions to deep networks can improve image classification accuracy compared to the corresponding control activation functions independently. Mixing activation functions across Swish and PTanh families had a larger benefit than within either family alone. We found larger improvements from mixed activation functions in a more constrained network. The dimensionality of the internal representation varied greatly depending on the choice of activation functions.

In this study we instantiated neural cell types by their activation functions, as a simplification of the *f-I* curves used to classify neurons, typically into “Type 1” and “Type 2” excitability. These types differ in how abruptly their *f-I* curves change in response to increase input current. Interestingly, one recent study found that “Type 1” and “Type 2” excitable neurons make spike timing networks more robust to correlated noise (Zeldenrust, 2021). Our findings suggest that diverse *f-I* curves may offer feed-forward computational benefits. However, some neurons exhibit nonlinear dynamics in their dendrites that are not effectively summarized by the *f-I* curve, underscoring that activation functions are an approximation of single neuron dynamics. Emerging work modeling dendritic computation (Beniaguev, 2019; Gidon, 2020) has parallels to network-in-network approaches in deep learning (Lin, 2013; Manessi, 2019). Beyond dynamical properties, there are many other attributes of neural diversity that are not commonly translated to deep learning, such as cell type specific sensory inputs, excitatory vs inhibitory neurons, and neuromodulation. Our findings join recent studies exploring the computational benefits of neural diversity, such as cell type specific connectivity (Stöckl, 2021), synaptic timescales (Burnham, 2021; Perez-Nieves, 2021), and membrane timescales (Perez-Nieves, 2021). Determining the functional role of neural cell types is an active area of research, and many of these cell type attributes may have translational benefit to deep learning applications.

An alternative approach to adding heterogeneity to network activation functions is to parameterize and learn the activation function parameters for each unit in the network. This parametric approach has been used with ReLU (He, 2015; Balaji, 2019), a set of basis functions (Goyal, 202), and piecewise linear functions (Agostinelli, 2015). Learning the activation function parameters has the benefit of increased flexibility at the cost of additional parameters, and choices about the parametric form of the activation functions. We find the largest benefit from mixing across parametric families of activation functions, demonstrating a distinct benefit beyond existing parametric approaches.

These findings raise an intriguing question: how do diverse nonlinearities impart improvements in feed-forward processing? One possibility is that mixed activation functions serve as more diverse basis functions within each layer, allowing increasingly complex transformations from one layer to another, increasing the overall expressivity of the network. A related possibility is that mixed activation functions balance the computational roles of sparsification and saturation across transformations of the network’s internal representation of the input space. Future work should further investigate the underlying mechanisms of diverse nonlinearities by examining the internal representations in mixed versus non-mixed networks. The relationship between network performance and internal representations is an active field of study, and mixed activation functions may serve as a useful test case for future work.

Our study explored the simplest alternating configurations for mixed activation functions in a 1:1 ratio. Ideally, automatic search techniques could generalize our approach to find optimal combinations of multiple activation function families across different layers of deep networks. Additionally, we found the highest performing activation functions in VGG11 and Sticknet8 differed. This highlights the need for further research to understand which activation functions, and in which combinations, lead to the largest improvement in accuracy. Our finding that a more constrained network had a larger benefit from mixed cell types suggests that one may see larger benefits on more complex datasets. In addition, future work should explore the role of mixed activation functions in settings beyond image classification. Deep learning is rapidly being applied to many tasks, and our study is only an initial investigation into the role of diverse activation functions.

In summary, heterogeneous cell types are a striking feature of biological circuits, and notably absent in deep artificial neural networks. In this study we instantiated diverse cell types by diverse activation functions. While many possible roles for cell types have been proposed, here we demonstrate their benefit in feed-forward computation in deep convolutional networks. This finding opens a large number of future studies to examine the role of cell types in biological circuits, and as tools in machine learning.

## Methods

### Network

Our base network architecture is VGG11 (Simonyan & Zisserman, 2014). The Pytorch instance of VGG11 contains 11 composite layers, 8 convolutional layers and 3 fully-connected layers. Each convolutional layer consists of a series of 3×3 convolutional filters, a ReLU activation layer, and a pooling layer that is included following activation layers 1, 2, 4, 6, and 8. Each fully-connected layer consists of a series of units linearly connected to input features with weights and biases, followed by a ReLU activation layer and a dropout layer. VGG11 contains 128 million trainable parameters.

Additionally, we developed a smaller network we named Sticknet8. Sticknet8 takes the first two convolutional layers from VGG and grafts them onto VGG’s final 3 fully-connected layers. Additionally, the number of units per layer decreased (Table 1). Sticknet8 contains 119,000 trainable parameters.

**Table 1.** Description of network architectures.

| Layer | VGG11 | Sticknet8 |
| --- | --- | --- |
| Conv1 | 64 filters | 8 filters |
| Activation1 | 64 units | 8 units |
| MaxPool1 | Kernel=2, stride=2 | Kernel=2, stride=2 |
| Conv2 | 128 filters | 16 filters |
| Activation2 | 128 units | 16 units |
| MaxPool2 | Kernel=2, stride=2 | Kernel=2, stride=2 |
| Conv3 | 256 filters | - |
| Activation3 | 256 units | - |
| Conv4 | 256 filters | - |
| Activation4 | 256 units | - |
| MaxPool4 | Kernel=2, stride=2 | - |
| Conv5 | 512 filters | - |
| Activation5 | 512 units | - |
| Conv6 | 512 filters | - |
| Activation6 | 512 units | - |
| MaxPool6 | Kernel=2, stride=2 | - |
| Conv7 | 512 filters | - |
| Activation7 | 512 units | - |
| Conv8 | 512 filters | - |
| Activation8 | 512 units | - |
| MaxPool8 | Kernel=2, stride=2 | - |
| FC9 | 4096 units | 128 units |
| Activation9 | 4096 units | 128 units |
| FC10 | 4096 units | 128 units |
| Activation10 | 4096 units | 128 units |
| FC11 | 4096 units | 128 units |

### Pytorch framework

We trained networks using Pytorch (Paszke, 2019). Each network had either the VGG11 or Sticknet8 architecture with random initial weights. Each network configuration is described by the names of their control nonlinearities and their corresponding parameter values. We initialized 10 instances of the base network with random weights, then replaced every ReLU layer with a MixedActivationLayer, alternating the control nonlinearities across each unit in the layer.

### Training dataset and procedure

We trained our networks for image classification on the CIFAR-10 dataset—a 10-class set of 32×32 pixel images with 6000 images per class, broken into 5000 training and 1000 validation images (Krizhevsky, 2009). Networks are optimized for top-1 accuracy via cross-entropy loss using the training algorithm Adam, which picks the best learning rate per network parameter (Kingma, 2015). This counteracts effects observed when using stochastic gradient descent where a global learning rate can lead to different observed rates of change in loss depending on the choice of activation function. For each network configuration, 10 random initializations are trained independently to assess statistical significance, using the same optimal learning rate selected for each network configuration. VGG11 and Sticknet8 networks were trained without using a learning rate scheduler for 500 or 300 epochs respectively.

**Figure 4.**
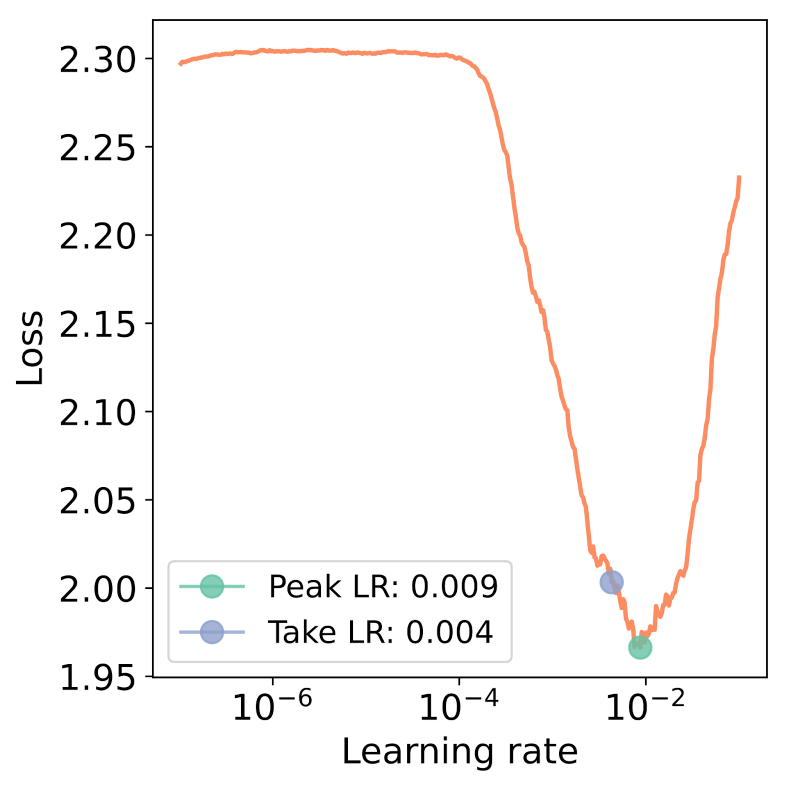
Learning rate vs. cross-entropy loss over a single training epoch. Lower learning rates fail to improve the loss, while higher learning rates cause it to diverge. The best value is somewhere on the negative slope between ~5e-4 and 1e-2 (Smith 2015). We initialized (Take learning rate, blue dot) each network configuration with 50% of the maximum learning rate (Peak learning rate, green dot).

### Network training and initial learning rate selection

Given that our networks contain mixtures of activation functions with different input/output scaling properties, it was important to use a training procedure that would optimize individual parameters at different learning rates. However, Adam - our optimizer (Kingma, 2015) - is sensitive to the initial global learning rate specified during instantiation. As with stochastic gradient descent, specifying too low an initial learning rate results in small updates to network parameters that fail to reduce the loss function over many training epochs, while specifying too high a learning rate prevents the optimizer from finding loss minima, leading to divergent behavior. This effect is particularly challenging in our case because a given network configuration can have its own optimum initial learning rate that if used to train other configurations can significantly hamper their learning, making comparisons of final accuracy less meaningful. Therefore we adopted a routine to choose the best initial learning rate per network configuration from a range of values based on loss recorded at each value (Smith 2015). The global learning rate is swept from low to high over the course of a single training epoch, stepping the learning rate and recording the loss after each minibatch. The best initial learning rate is then determined to be the highest value before loss starts to diverge, i.e. the lowest point of the LR-loss profile in Figure 4. This value is averaged across 10 network initializations per configuration, divided by 2 to account for variance in the LR-loss profile across initializations that might cause the routine to select too high a value, and given as input to the Adam optimizer at the start of training.

### Max and Linear Predictions

For each mixed network, we define a maximum and linear prediction. For each corresponding control network, we compute the average validation accuracy across the 10 random initiations, and refer to this as the control accuracy. The maximum prediction is the maximum control accuracy across all corresponding control networks. The linear prediction is the weighted average between the corresponding control networks. Unless otherwise specified, our control networks were mixed in a 1:1 ratio, so the weighted average was a simple average. Where presented in the text, 95% confidence intervals were computed by assuming normally distributed variations across the 10 random initiations.

To establish statistical significance for our mixed networks compared to their control cases we utilized a 1-sided t-test between control and mixed networks. Pairs of initializations from both groups were randomly selected. We corrected for multiple comparisons using the Benjamini-Hochberg correction to produce a corrected false-positive rate of p=0.05.

## Appendix

The full code repository, containing all training, analysis, and visualization code can be found here: https://github.com/briardoty/allen-inst-cell-types

